# Human Microbiota of the Argentine Population- A pilot study

**DOI:** 10.1101/030361

**Authors:** Belén Carbonetto, Mónica Carolina Fabbro, Mariela Sciara, Analía Seravalle, Guadalupe Méjico, Santiago Revale, María Soledad Romero, Bianca Brun, Marcelo Fay, Fabián Fay, Martin Vazquez

## Abstract

Here we present the first dataset based on human microbiota samples of an urban middle-income population in South America. We characterized the microbiota of six different body habitats: palatine tonsils, saliva, buccal mucosa, throat, anterior nares and gut from samples of healthy individual living in a metropolitan area in Argentina. Our initial findings revealed differences in the structure and composition of the microbial communities compared to the US urban population. By sharing our data, we want to actively encourage its reuse for comparison purposes. This  will hopefully result in novel biological insights on the variability of the microbiota of healthy individuals across populations worldwide. Moreover, the understanding of the human microbiota ecosystem in a health-associated state will help to answer questions related to the role of the microbiota in disease.

## Background

The human microbiota is the collection of microorganisms living in or on the human body. An imbalance or dysbiosis in these microbial communities can be associated with disease (Pham and Lawley, 2014, Petersen and Round, 2014, Zaura et al., 2014). Moreover, when the microbiota of the same bodysites are compared between different healthy individuals, specific microbial community features are apparent (Relman, 2015, Li et al., 2012, Zhou et al., 2013). In addition differing selective pressures are found at distinct body sites leading to different patterns in microbial community structure and composition (Yatsunenko et al., 2012, Oh et al., 2014, Stearns et al., 2011). Because of the existence of these natural variations, a comprehensive characterization of the healthy microbiota is critical for predicting alterations related to disease. This characterization should be based on a broad population of healthy humans over time, geography and cultural traditions (Yatsunenko et al., 2012, Shetty et al., 2013, Ross et al., 2015, Leung et al., 2015). The study of healthy individuals representing different ages, cultural traditions and ethnic origins will enable to understand how the associated microbiota varies between populations and respond to different lifestyles. It is important to address these natural variations in order to later detect other kind of variations that are related to disease.

During the last decade researchers from around the world have characterized and defined geographical differences in the composition of microbiota in healthy adult humans. The most important projects so far have been the Human Microbiome Project (HMP) (Peterson et al., 2009), MetaHIT (Arumugam et al., 2011, Qin et al., 2010) and the American Gut Project. Despite the big effort conducted, these major projects are based on American and European populations and have not covered all ethnic groups and different socio-economic, geographic and cultural settings. Some projects have been conducted to feel this gap (Yatsunenko et al., 2012, Ross et al., 2015, Clemente et al., 2015, Leung et al., 2015). Nevertheless there are still no records on human microbiota of urban middle-income populations in South America. Here we characterized the microbiota of six different body habitats: palatine tonsils, saliva, buccal mucosa, throat, anterior nares and gut from samples of 20 healthy middle-income male and females between the ages of 20 and 50 years living in Rosario City, Argentina. Genomic DNA was purified from each sample and the 16S rRNA gene V1-V3 region was amplified. Amplicons were sequenced on a 454 GS-FLX+ Titanium platform and produced a total of 323,110 filtered reads. We compared microbial community composition and structure data with the same data derived from the Human Microbiome Project. Results showed that microbiota of buccal mucosa, palatine tonsils and gut microbiota differed between the Argentine and American populations. We made publically available the first 16S rDNA profile dataset of human body microbiota of an Argentine urban cohort. Our results enhance the idea that the generation of more and larger local datasets of healthy individuals is needed in order to analyze different microbiota dysbiosis related to disease in the Argentine middle-income urban population.

## Methods

### Sampling

Samples were collected in a non-invasive manner from palatine tonsils, saliva, buccal mucosa, throat, anterior nares and gastrointestinal tract (stool) of healthy men and women between 20 and 50 years old, who live in Rosario city in the central region of Argentina. Subjects donated blood to examine the presence of viral markers and metadata was collected by medical examination in order to select healthy individuals (Supp. Table1). Samples were collected in a single visit and sent immediately to the lab where they were processed.

### Ethics, consent and permissions

The study was approved by the Institutional Ethic’s Committee from the Hospital Italiano Garibaldi in Rosario, Argentina. Moreover, a consent form with information about the study, including the rights of the subject and the risks and benefits involved in participating in the study was signed by each individual.

### DNA extraction and sequencing

Total genomic DNA was extracted from 200mg of each fresh stool sample using QIAamp (Qiagen, CA) DNA Stool Mini Kit following manufacturer’s instructions. Palatine tonsils, saliva, buccal mucosa, throat and anterior nares swabs were first resuspended in 200μl steril saline solution. Genome DNA was extracted from this solution using QIAamp (Qiagen, CA) DNA Mini Kit following manufacturer’s instructions.

For the construction of pyrotag libraries the V1-V3 hyper variable regions of the 16s rRNA gene was amplified using the 27F (5’AGAGTTTGATCCTGGCTCAG3’) and 534R (5’ATTACCGCGGCTGCTGG3’) tagged primers (2012). Samples were amplified using two rounds of PCR, the first to amplify the 16S rRNA gene (30 cycles) and a second short round to add barcodes for sample identification (10 cycles), following the procedures detailed in Rascovan et al. 2013 (Rascovan et al., 2013). Duplicated reactions were performed in both rounds of PCR to reduce amplification biases and then pooled. All amplicons were cleaned using Ampure DNA capture beads (Agencourt-Beckman Coulter, Inc.) and pooled in equimolar concentrations before sequencing on a Genome Sequencer FLX (454-Roche Applied Sciences) using 454 GS FLX+ chemistry according to the manufacturer’s instructions.

### Amplicon sequence processing, OTU classification and taxonomic assignment

Our dataset and the HMP dataset (http://hmpdacc.org/HMQCP/) were processed using the QIIME v1.8 analysis pipeline (Caporaso et al., 2010b). For comparative purposes the same number of individuals (N=20) was selected randomly from the HMP dataset. First a random selection of 1000 reads per sample was done using a custom-made script. Sequences were then clustered into Operational Taxonomic Units (OTUs) using the pick_otus.py script with the Uclust method at 97% sequence similarity (Edgar, 2010). OTU representative sequences were aligned using PyNast algorithm with QIIME default parameters (Caporaso et al., 2010a). Phylogenetic trees containing the aligned sequences were then produced using FastTree (Price et al., 2009). Richness alpha diversity metrics and rarefaction curves were calculated by sub-sampling the OTU tables at different depths and counting the resulting number of phylotypes using 10 iterations per sample. Phylogeny-based beta diversity distances between OTUs were calculated using weighted Unifrac (Lozupone and Knight, 2005, Lozupone et al., 2007). Taxonomic classification of sequences was done with Ribosomal Database Project (RDP) Classifier using the Greengenes V13.5 database and a 50% confidence threshold (Wang et al., 2007).

### Numerical analyses

Unifrac phylogenetic pairwise distances among samples were visualized with principal coordinates analysis. Analysis of similarity statistics was calculated to test a-priori sampling groups. Mann-Whitney non-parametric tests were performed to elucidate differences in taxa abundances. All calculations were carried out with R packages ‘BiodiversityR’ and ‘Vegan’.

## Results

### Argentine human microbiota overview

Results showed that OTU richness differed between body habitats. Saliva presented the higher richness and the nose was the less diverse site (Figure 1A). Beta diversity also revealed differences in community structure between habitats (Figure 1B). The PCoA visualization revealed a separation of the data in three main groups: oropharyngeal region, nose and gut (ANOSIM p<0.05).

**Figure 1:**
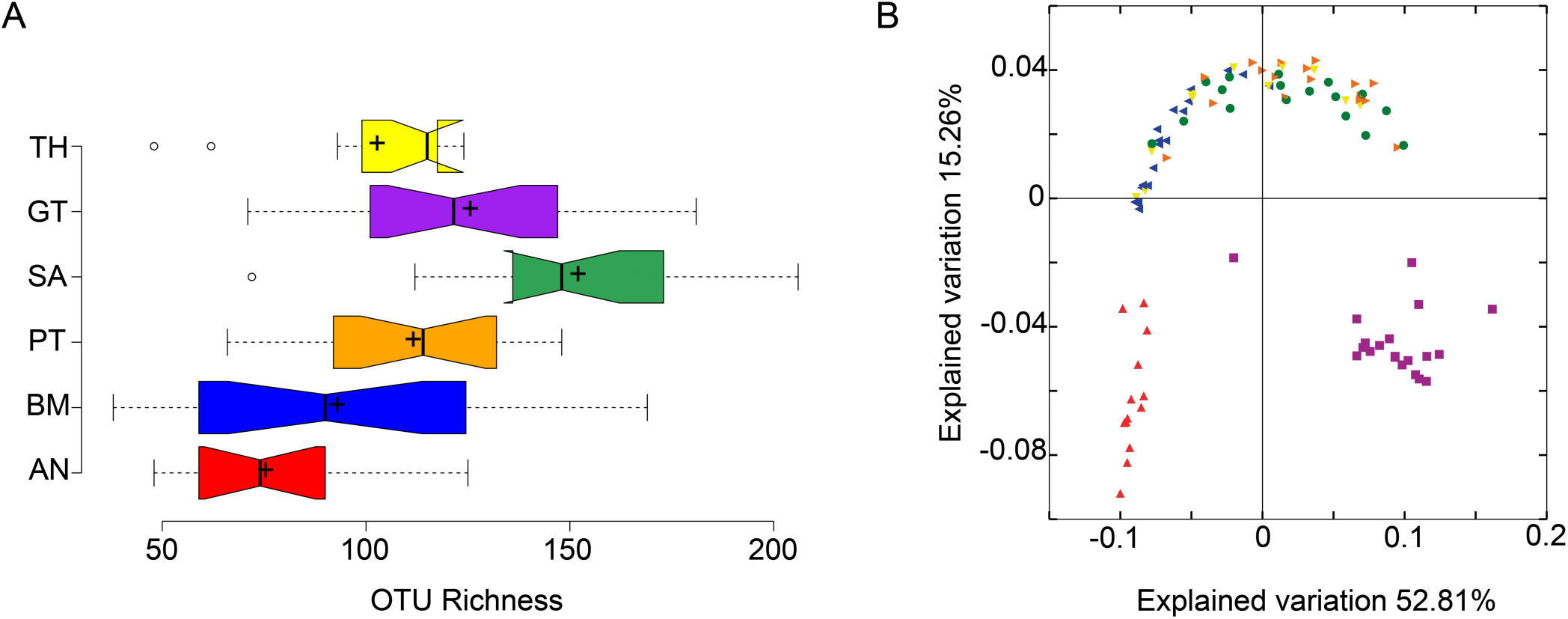
Argentine human microbiota diversity analysis. (A) Alpha diversity analysis based on OTU richness BM stands for buccal mucosa, TH for throat, PT for palatine tonsils, AN for anterior nares, SA for saliva and GT for gut. (B).Beta diversity analysis based on Weighted unifrac pairwise distances, each color represent a body site as in A.

### Comparison with the HMP dataset

We observed that the microbiota of the Argentine and American populations differed in composition and community structure. Beta diversity results based on Weighted Unifrac distances showed that the microbiota of the buccal mucosa, the palatine tonsils and the gastrointestinal tract differed between Argentine and American individuals (Supp. Figure 1, ANOSIM p<0.05). Moreover, the taxonomic composition of these body habitats was also different between populations. We observed differences in the abundance of the most predominant taxa (Figure 2). For example the abundance of Bacteroidaceae family was higher in the American population gut microbiota, while Ruminococcaceae, Lachnospiraceae, Rikenellaceae and Prevotellaceae were more abundant in the Argentine gut microbiota. It is known that the variation in the levels of three main taxa in the gut microbiota can define enterotypes (Arumugam et al., 2011). The higher relative abundance of three genera defines the enterotypes: Bacteroides defines enterotype 1, Prevotella defines enterotype 2 and of Ruminococcus defines enterotype 3. Although we found differences in Bacteroidaceae and Ruminococcaceae abundances (Figure 2, Mann-Withney p<0.05) and Bacteroides, Ruminococcus and Prevotella abundances (Mann-Withney p<0.05, data not shown) between populations, both populations can be assigned to enterotype 1 (Supp. Figure 2). This is, to our knowledge the first time gut microbiota of middle-income populations of North and South America are compared. Other author’s did found differences in Prevotella and Bacteroides abundances between southamerican and US populations, but they compared ameridian individuals living in little villages near the Amazonas with western citizens living in metropolitan areas (Yatsunenko et al., 2012). Southamerican ameridians presented higher Prevotella abundance than US individuals. As expected the authors hypothesized that differences in diet were mainly explaining differences in the gut microbiota. Our results encourage this hyphothesis. We observed differences in the relative abundances of the most important taxa but the Bacteroides/Prevotella trade-off (enterotype 1) was the same for the argentine and US populations. The diet of people in the main metropolitan areas of Argentina is more similar to the one of individuals in the main cities of the US than it is from individuals in the mentioned villages near the Amazonas, which is mainly based on maize and native roots rather than processed food. Our initial findings support this idea that diet is more relevant when defining gut enterotypes than it it is geographic location or ethnics. Nevertheless, the difference in taxa abundances between our dataset and the HMP dataset reflects the differences in cultural traditions in the Argentine and US cities. As regards the buccal mucosa microbiota we observed that Streptococcaceae family was more abundant in the Argentine population while Gemellaceae family was higher in the US individuals (Figure 2, Mann-Withney p<0.05). The abundance of Streptococcaceae family was also higher in the palatine tonsils of argentine individuals. We hypothesize that the differences in the observed oral/pharyngeal profiles may be due to differences in the use of antibiotic treatments in both populations. Long-term impacts in oral microbiota due to antibiotic administration has been reported, moreover antibiotic resistance were observed in these cases (Jakobsson et al., 2010). Although this is just a hypothesis that needs to be proven our results encourage the idea that the human microbiota ecosystem has multiple states of equilibrium and that these variations are present between and within healthy populations. Then, these multiple states related to different lifestyles, location, ethnics, cultural tradition, age, gender, etc. need to be studied in depth in order to further understand alterations on the equilibrium due to disease.

**Figure 2:**
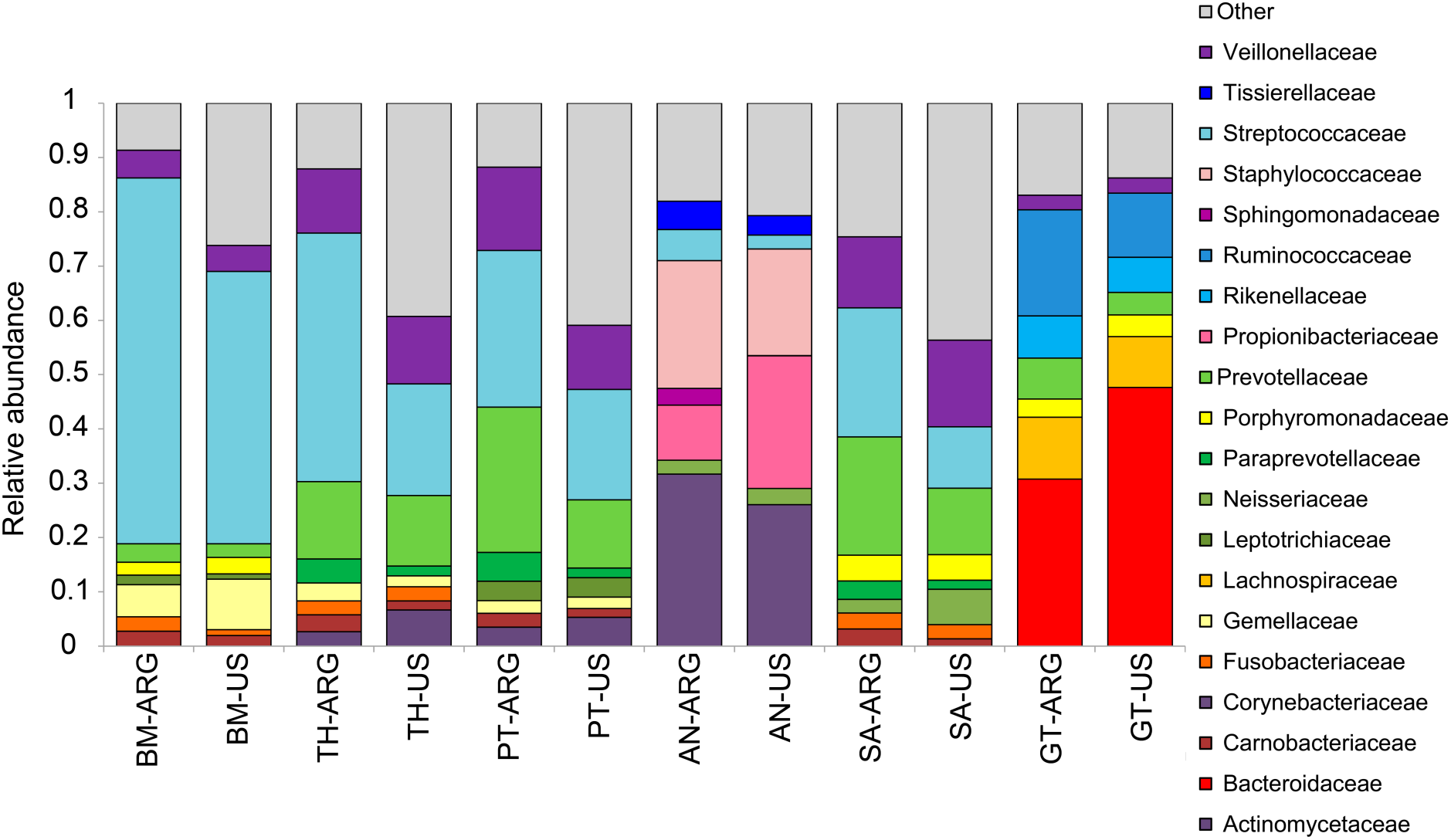
Taxonomic profiles. Most abundant bacterial families in the Argentine and American (HMP) human microbiota. ARG refers to the Argentina population and US to the American population. BM stands for buccal mucosa, TH for throat, PT for palatine tonsils, AN for anterior nares, SA for saliva and GT for gut.

## Conclusions

Here we present the first dataset based on human microbiota samples of an urban middle-income population in South America. We characterized the microbiota of six different body habitats: palatine tonsils, saliva, buccal mucosa, throat, anterior nares and gut from samples of healthy individual living in a metropolitan area in Argentina. Our initial findings revealed differences in the structure and composition of the microbial communities compared to the US urban population.

By sharing our data, we want to actively encourage its reuse for comparison purposes. This will hopefully result in novel biological insights on the variability of the microbiota of healthy individuals across populations worldwide. Moreover, the understanding of the human microbiota ecosystem in a health-associated state will help to answer questions related to the role of the microbiota in disease.

## Data Access

Datasets supporting the results of this study are available in the NCBI SRA database under the BioProject with accession number: PRJNA293521 (http://www.ncbi.nlm.nih.gov/bioproject/PRJNA293521/).

## Conflict or interest statement

The authors declare that they have no conflict of interest.

## Authors’ contributions

BC and MCF analyzed the data and performed the numerical and statistical analyses. BC wrote the manuscript. SR processed the sequence raw data. MF and MS designed the protocol and criteria for healthy individual selection. MS wrote the protocol for recruiting the individuals, supervised the recruiting process, and collected the metadata for the individuals. GM performed DNA preparation from human samples. MSR and BB performed amplicon library preparation and sequencing. FF directed the project and supervised the study. MV directed the project, conceived the study, supervised the bioinformatics analysis and supervised the manuscript.

